# MicroCT enables simultaneous longitudinal tracking of murine pancreatic cancer progression and cachexia

**DOI:** 10.1101/2025.05.29.656943

**Authors:** Katherine R. Pelz, Philip Jimenez, Colin J. Daniel, Samuel D. Newton, Melissa Cunningham, Rosalie C. Sears, Patrick J. Worth, Jonathan R. Brody, Teresa A. Zimmers

**Author notes:** Corresponding author: Teresa A. Zimmers, PhD, Department of Cell, Developmental and Cancer Biology, School of Medicine, Oregon Health & Science University, Portland, OR.

## Abstract

Preclinical models of pancreatic ductal adenocarcinoma (PDAC) can greatly benefit from non-invasive imaging for evaluating disease progression and therapeutic response. Imaging approaches that can accurately and simultaneously track primary tumor growth, metastatic dissemination, and host cachexia over time are lacking. Here, we report an optimized dual-contrast micro-computed tomography (microCT) protocol for longitudinal imaging in orthotopic murine models of PDAC. This method enables high-resolution, volumetric quantification of orthotopic primary tumors, liver and lung metastases, and paraspinal skeletal muscle, providing a dynamic view of both tumor and host physiology. MicroCT primary tumor measurements strongly correlated with endpoint tumor weights and outperformed 2D ultrasound in early detection and volumetric accuracy, particularly for small or irregularly shaped tumors. This platform revealed heterogeneous metastatic kinetics across PDAC models and uncovered early, heterogeneous onset of skeletal muscle wasting, a hallmark of cancer cachexia. Notably, this protocol mimics clinical CT surveillance by enabling opportunistic cachexia assessment from tumor imaging datasets and offers substantial advantages over destructive endpoint analyses. Further, microCT radiation had no effect on our model endpoints. By capturing the temporal dynamics of tumor progression and host response, dual contrast microCT serves as a powerful translational platform for preclinical PDAC research and therapeutic testing.

**Significance:** Dual-contrast microCT provides high-resolution, whole-body, non-invasive imaging in orthotopic murine PDAC models, enabling simultaneous tracking of tumor growth, metastasis, and skeletal muscle wasting–offering a clinically relevant, translational imaging platform.

## Introduction

Pancreatic ductal adenocarcinoma (PDAC) remains one of the deadliest malignancies, with poor prognosis driven by treatment resistance, aggressive metastatic spread, and high prevalence and severity of cancer-associated cachexia^1–3^. Pre-clinical studies of PDAC rely heavily upon genetic and orthotopic mouse models^4,5^, however longitudinal monitoring of PDAC progression—including primary tumor growth, the timing of metastasis, and associated cachexia—remains challenging. Common imaging modalities used in murine PDAC models include bioluminescence imaging (BLI) and ultrasound (US)^6^. BLI provides a convenient readout of tumor burden, but lacks spatial resolution, limiting its utility for accurate volumetric measurements and organ-specific detection of metastases^7^. BLI provides no information about body composition, obviating use for studies of cachexia. US offers higher resolution, avoids radiation exposure, and can be used for muscle measurements to assess cachexia severity. However, conventional two-dimensional (2D) US often assumes spherical tumor geometry, reducing accuracy. While 3D US improves volumetric accuracy for primary tumor measurements^8^, effectiveness of US in detecting metastases is limited due to rib interference when imaging common metastatic sites like the liver and lungs. Together these limitations underscore the need for a robust, high-resolution imaging approach capable of providing detailed longitudinal measurements of both tumor progression and host wasting.

To address this gap, we hypothesized that micro-computed tomography (microCT) imaging in conjunction with contrast administration could be used to track pancreatic cancer progression longitudinally in murine models, providing insights into the temporal dynamics of primary tumor growth, metastatic dissemination, and cachexia. MicroCT offers high spatial resolution and the capacity for whole-body imaging, making it a promising tool for comprehensive tumor monitoring (Supplementary Fig. 1)^9^. However, to our knowledge, there are no protocols available describing the use of microCT for concurrent monitoring of PDAC primary tumor, metastases, and cachexia.

Herein, we report an optimized dual-contrast microCT protocol for serial, non-invasive imaging of orthotopic pancreatic tumors. This method enables simultaneous high-resolution tracking of primary tumor growth, metastasis, and concurrent muscle wasting, providing precise, longitudinal data on PDAC and host dynamics with minimal impact on animal health.

## Results

### MicroCT enables accurate primary tumor volume quantification and growth kinetics in murine PDAC orthotopic models

To evaluate the sensitivity of microCT for preclinical imaging, we used a murine PDAC model with orthotopic injection of 5,000 pancreatic tumor cells or PBS (control), monitoring animals until they reached moribund status per IACUC guidelines (Hickman score ≤ 3^10^ or tumor weight ≥ 10% body weight) (**Fig. 1**). We tested four PDAC cell lines with distinct phenotypes and metastatic propensities: KPC 32908, KPC 7107, KPC 8060, and KMC Z682. KPC were derived from LSL-*Kras^G12D/+^;*LSL-*Tp53^R172H/+^;Pdx1-Cre* mice^11^ and KMC from LSL-*Kras^G12D/+^*;*Rosa26*-LSL*^Myc/Myc^*;*Ptf1a^CreERTM^*mice^12^. Scans began one-week post-injection to allow for surgical recovery, re-scanning every 3–4 days (**Fig. 1**). Primary endpoints included longitudinal primary tumor volume, liver and lung metastasis volume, skeletal muscle area, and latency to metastasis.

**Figure 1.**
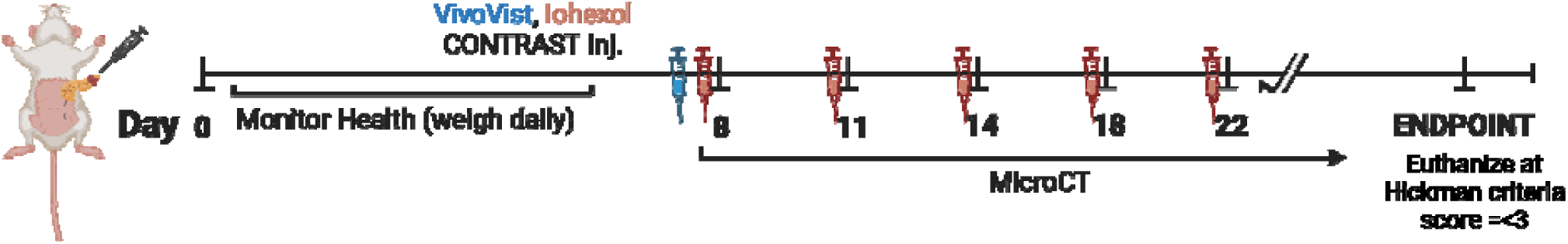
Study experimental timeline. Mice were orthotopically implanted with PDAC cells or PBS (day 0), injected with Vivovist (24 hours before the first scan) and iohexol (before each scan), and scanned every 3-4 days starting on day 8. Mice were euthanized upon reaching humane criteria (Hickman score ≤ 3^10^).

To visualize PDAC tumors, iohexol (300 mg I/mL) diluted 1:3 in PBS (final concentration of 1.2 mg iodine per gram of body weight) was injected intraperitoneally (IP) immediately prior to each scan (**Fig. 1**), providing bright contrast throughout the peritoneal cavity to delineate the primary tumor from surrounding organs (**Fig. 2a**, Supplementary Fig. 2). Iohexol at this dose was metabolized within 20 minutes (Supplementary Fig. 3). Using iohexol contrast with microCT imaging, we accurately quantified primary tumor volumes in our orthotopic PDAC model and generated 3D tumor images (**Fig. 2b**). MicroCT-derived volumes strongly correlated with endpoint tumor weights (R² = 0.9475, p < 0.0001, **Fig. 2c**). Additionally, we compared microCT volumes to measurements obtained from 2D ultrasound (US) using axial and sagittal areas with x-, y-, and z-coordinates (**Fig. 2d**). While there was a positive correlation between US-and microCT-derived volumes overall (R² = 0.6349, p = 0.0011, **Fig. 2e**), the correlation was lost for small tumors (< 700 mm^3^ by microCT) (R² = 0.00, p = 0.9839, **Fig. 2f**), and US consistently underestimated volumes, likely due to non-spherical tumor shape. These findings highlight microCT as a superior imaging modality for accurate primary tumor measurement, particularly during early tumor progression. The accuracy of microCT tumor measurements was further demonstrated in subcutaneous C26 carcinoma models (Supplementary Fig. 4).

**Figure 2.**
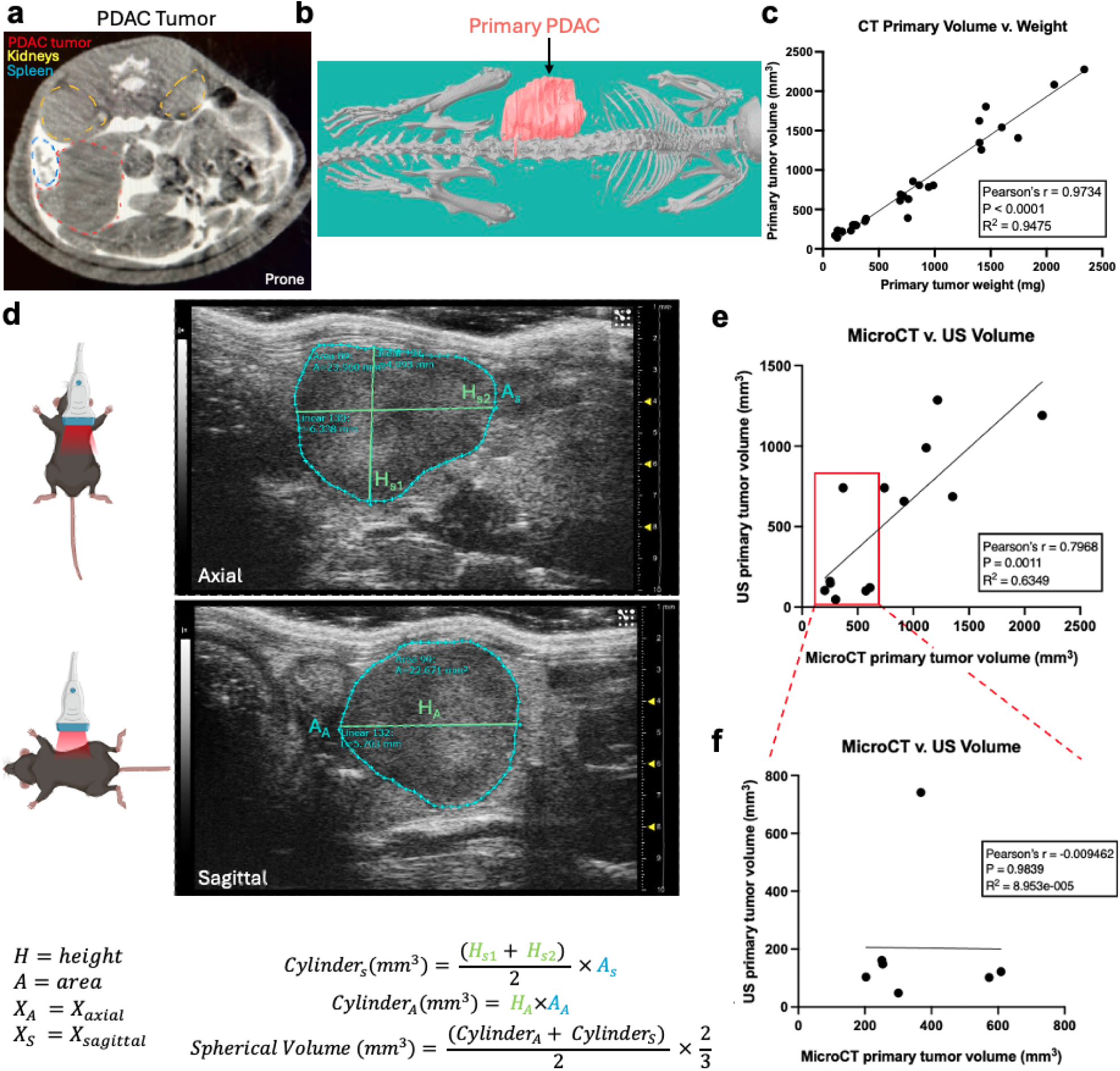
Longitudinal monitoring of PDAC tumor kinetics using microCT imaging and volumetric comparisons with 2D ultrasound (US). **a)** Representative axial microCT image showing primary tumor (red outline). **b)** 3D reconstruction showing the primary tumor (red) and skeleton of the mouse rendered in grayscale. **c)** Correlation of primary tumor volume measured by microCT with tumor weight at endpoint (N = 27, Pearson’s r = 0.9734, p < 0.0001, R² = 0.9475). **d)** Schematic and representative US images showing how primary tumor volume was estimated by 2D US. Axial and sagittal slices at the largest tumor dimensions were obtained, and area (A) and height (H) measurements were used to derive cylindrical volumes for each plane. Tumor volumes were calculated by averaging the volumes of the axial and sagittal cylinders and applying a spherical volume correction factor (2/3). **e)** Correlation between US- and microCT-derived primary tumor volumes (R² = 0.6349, p = 0.0011). **f)** Correlation between US- and microCT-derived primary tumor volumes for subset of tumors smaller than 700 mm³ (as determined by microCT) (R² = 0.00, p = 0.9839).

### MicroCT allows non-invasive, high-resolution tracking of PDAC liver and lung metastases

To visualize PDAC liver metastases, a single IP injection of 1.5 g/kg Vivovist (Nanoprobes #1301) was administered 24 hours before the first scan (**Fig. 1**), which enhanced contrast between liver tissue and metastases (**Fig. 3a**). Vivovist, composed of alkaline earth metal nanoparticles, is rapidly taken up by the liver and spleen, maintaining consistent contrast intensity for the study duration (up to 72 days post-injection) (Supplementary Figs. 5–7). Lung metastases require no additional contrast, as air provides sufficient differentiation (**Fig. 3b**). A limitation of this protocol is the persistent contrast artifact from Vivovist in liver and spleen histopathology, presenting as a dark brown hue on both hematoxylin and eosin (H&E) and immunohistochemistry (IHC) stains (Supplementary Fig. 8). To mitigate this, we recommend immunofluorescence for protein analysis or alkaline-phosphatase-conjugated antibodies for IHC, as red staining is distinguishable from brown artifact when analyzed with software such as ImageJ (Supplementary Fig. 8).

**Figure 3.**
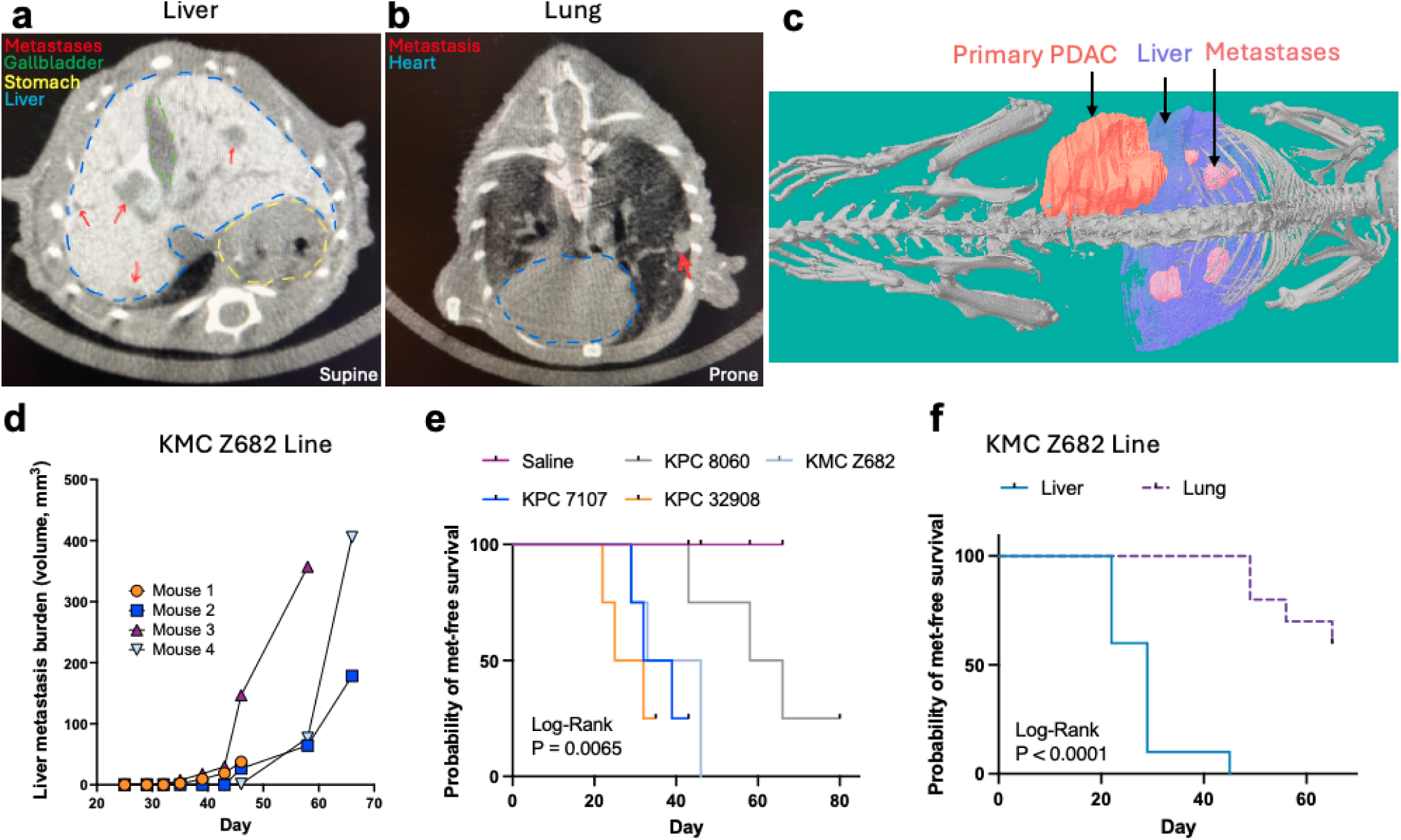
Longitudinal monitoring of PDAC liver and lung metastasis using microCT imaging. a-b) Representative axial microCT images showing **a)** liver metastases (red arrows) and **b)** lung metastasis (red arrow). **c)** 3D reconstruction showing the primary tumor (red), liver (blue), metastases (pink), and skeleton of the mouse rendered in grayscale. **d)** Longitudinal live metastatic burden in KMC Z682-injected mice (N = 4). **e)** Metastasis-free survival for PDAC lines (N = 4/group, p = 0.0065). **f)** Liver versus lung-specific metastasis-free survival in KMC Z682-injected mice (N = 10 mice, Log-Rank p < 0.0001).

MicroCT imaging with Vivovist contrast enabled precise, non-invasive tracking of metastasis progression (**Fig. 3c**). We focused on liver and lung metastases, the most common sites of metastasis in PDAC^13^, however metastases were also detected in the spleen in images and at euthanasia (Supplementary Fig. 9). Longitudinal monitoring allowed volumetric quantification of liver metastases over time (**Fig. 3d**). Our orthotopic PDAC models exhibited substantial inter- and intra-line heterogeneity, emphasizing the importance of serial imaging for robust comparisons. Longitudinal analyses improve sensitivity over single time-point comparisons, enhancing statistical power through repeated-measures models rather than t-tests at one time-point. Kaplan-Meier analysis showed distinct metastatic timing among PDAC lines: KPC 32908 (28.5 days), KPC 7107 (35.5 days), KMC Z682 (39.5 days), and KPC 8060 (62 days) (p = 0.0065, log-rank test, **Fig. 3e**). In the KMC Z682 model, mice frequently develop lung metastases in addition to liver metastases. MicroCT identified emergence of liver metastases significantly earlier than lung metastases in this line (median 29 vs. 52.5 days, p < 0.0001, log-rank test, **Fig. 3f**). Compared to US, microCT provided superior consistency and spatial resolution when tracking liver and lung metastases in longitudinal studies (Supplementary Table 1). The ability to quantify metastatic burden volumetrically via microCT also offers distinct advantages over endpoint histological or flow cytometry-based analyses, which require large cohort sizes and extensive tissue processing. Histology and flow cytometry demand complete serial sectioning or tissue digestion to detect small lesions, whereas microCT captured whole-organ metastasis non-invasively.

### Longitudinal microCT imaging reveals early and heterogenous onset of cachexia-induced skeletal muscle wasting

Analogous to opportunistic use of clinical diagnostic CT scans to quantify muscle loss–the current gold standard for monitoring cachexia in clinical studies—we tracked paraspinal muscle area longitudinally (**Fig. 4a,b**). All PDAC lines induced significant muscle wasting. KPC 32908 produced the most rapid decline; muscle area decreased by 43% by day 29, when all other lines were reduced by <10% (**Fig. 4c**). Significant decreases in paraspinal muscle area were first detected at d29 and d35 for KPC 32908 and 7107, respectively (Dunnett’s multiple comparisons test, P=0.0210, P=0.0188). However, paraspinal muscle weights showed a downward trend as early as days 18, 25, 35, and 39 for KPC 32908, KPC 7107, KMC Z682, and KPC 8060, respectively (**Fig. 4c**). All PDAC models showed signs of cachexia at endpoint by standard measures– hindlimb muscle weights normalized to initial body weight (30–33% loss) (**Fig. 4d**). The KPC 32908, KPC 7107, and KMC Z682 groups also had significantly reduced heart weights compared to controls (Supplementary Fig. 10). These findings highlight both the prevalence of cachexia in PDAC, and that skeletal muscle wasting is an early manifestation of the disease. Notably, different PDAC models demonstrated variable cachexia-inducing capacities at early time points. This heterogeneity would be challenging to detect using endpoint analyses alone, highlighting the importance of longitudinal microCT imaging for capturing dynamic changes in muscle wasting and better understanding the progression of cachexia in preclinical cancer models.

**Figure 4.**
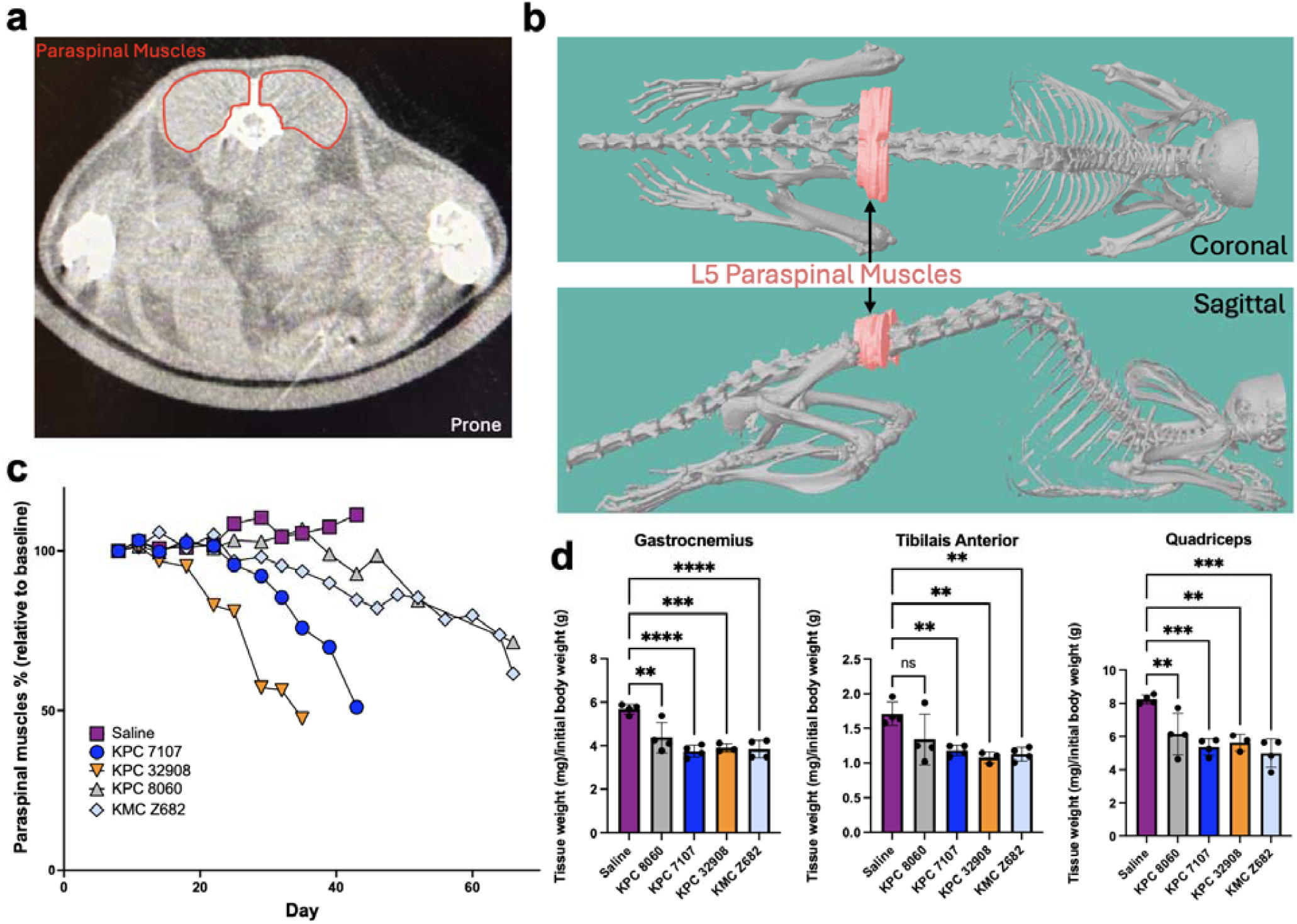
Longitudinal assessment of skeletal muscle wasting by microCT. **a)** Representative axial microCT image showing paraspinal muscle cross-section at the L5 vertebra (red outline) —the region used for area measurements. **b)** 3D reconstruction of microCT scan showing L5 paraspinal muscles (red) in the coronal and sagittal planes. **c)** Longitudinal tracking of paraspinal muscle area, expressed as a percentage of baseline, for mice injected with different PDAC lines or PBS control. **d)** Post-mortem analysis of gastrocnemius, tibialis anterior, and quadriceps muscle weights normalized to initial body weight across experimental groups (N = 4 per group). Statistical analysis used one-way ANOVA with Dunnett’s multiple comparisons test. ns = not significant, ** p < 0.01, *** p < 0.001, **** p < 0.0001. Error bars represent mean ± standard deviation.

### Imaging frequency considerations to balance data resolution with radiation exposure

Of note, this approach parallels clinical CT imaging used to track tumor progression in patients, where scans typically occur every three months^14^. Given the short lifespan of mice, our imaging frequency provides a comparable timescale for translational relevance. However, frequent scanning resulted in substantial radiation exposure for the mice (mean cumulative dose per mouse = 10 Gy over 66 days, Supplementary Table 2) and investigator time (4 minutes per scan and ∼15 minutes for reconstruction and analysis). Therefore, we recommend only using this scan frequency for initial cell line or model characterization, with reduced imaging intervals tailored to desired experimental endpoints in subsequent studies. Although we did not observe overt radiation-induced effects on mouse health, distinguishing between radiation-specific and tumor-driven effects was challenging due to overlapping sickness behaviors. Comparisons between PBS control mice (receiving the same radiation regimen but without tumors) and PDAC-bearing mice indicated that the observed weight loss and muscle wasting were tumor-induced (**Fig. 4d**, Supplementary Fig. 11). To confirm that radiation had no effect on our model endpoints, we performed a study following our standard protocol (**Fig. 1**) but included a matched group that received contrast injections and anesthesia without microCT scanning (**Fig. 5a**). We found no differences in primary tumor weight or primary tumor and liver metastases volumes measured by microCT at euthanasia (day 43 post-injection) between groups (**Fig. 5b-d**), confirming that the radiation doses proposed in our protocol do not impact our model endpoints. Body weight trajectories were comparable between groups (**Fig. 5e**), as were skeletal muscle, heart, and epididymal adipose tissue weights at euthanasia (**Fig. 5f-i**). Radiation reduced testis size, consistent with previous reports^15^ (**Fig. 5j**). Regardless, careful consideration of cumulative radiation exposure is essential, and scan frequency should be optimized to balance data collection needs and minimize radiation-related confounders. Indeed, in subsequent studies using the KMC Z682 line, we have scanned weekly from day 15, increasing to every 3 days between days 29–46 to capture time-to-metastasis, and reverting to weekly scans thereafter to track tumor growth, metastasis volume, and skeletal muscle area.

**Figure 5.**
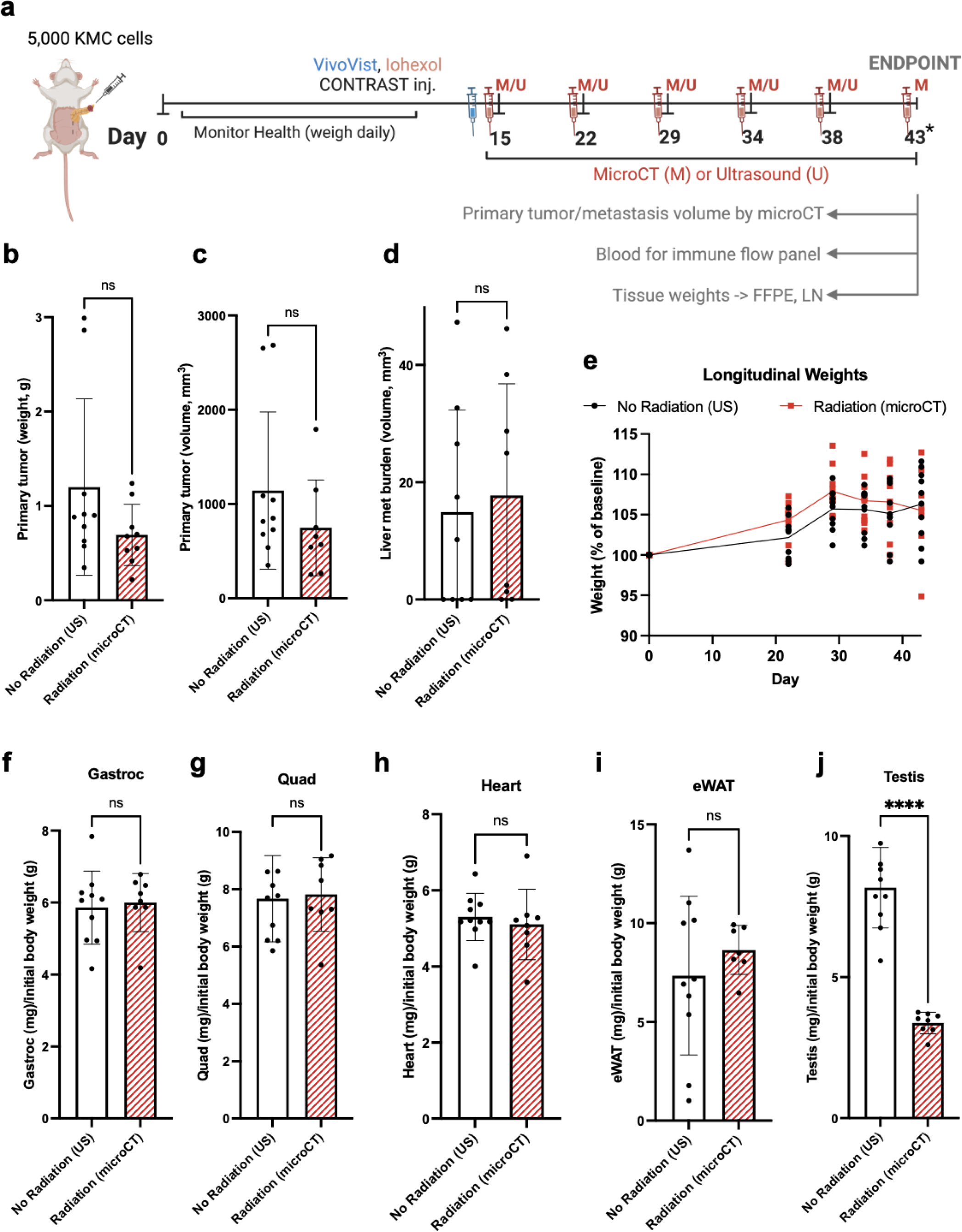
Radiation exposure effects on PDAC model tumorigenesis and muscle/adipose tissue weights. **a)** Experimental timeline: Mice were injected orthotopically with 5,000 KMC Z682 cells and longitudinal imaging was performed via microCT (M) or ultrasound (U) at specified time points. All mice underwent microCT scanning immediately prior to euthanasia. Average radiation dose per scan was 715 mGy. **b)** Primary tumor weight (g) and **c)** primary tumor and **d)** liver metastasis volume (mm³) at euthanasia. **e)** Longitudinal body weight represented as % of baseline. **f-j)** Tissue weights normalized to initial body weight at euthanasia. **f)** Gastrocnemius, **g)** quadriceps, **h)** heart, **i)** epididymal white adipose tissue, and **j)** testis. Statistical analysis was performed using t-tests. **** p < 0.0001, ns = not significant. Error bars represent mean ± standard deviation.

## Discussion

In this study, we developed and validated a dual-contrast microCT protocol for the preclinical monitoring of pancreatic cancer progression in murine models. By providing simultaneous, precise, longitudinal volumetric measurements of tumors, metastases to different organ sites, and cachexia-associated skeletal muscle wasting, our protocol surpasses conventional imaging methods like BLI and US (Supplementary Table 1), which often lack spatial resolution, suffer from signal attenuation due to tissue depth, or fail to simultaneously quantify tumor burden and host tissue changes.

To our knowledge, this is the first study to describe the dual use of iohexol and Vivovist contrast agents for the simultaneous delineation of primary pancreatic tumors and liver metastases via microCT. Consistent with prior work, we found that intraperitoneal administration of iohexol effectively enhances soft tissue contrast in the peritoneal cavity, enabling precise localization of PDAC tumors and improved delineation from surrounding organs^16,17^. While previous studies used fixed volumes of iohexol ranging from 250 μL to 3 mL, we found that injecting volumes closer to 100 μl of iohexol diluted 1:3 in PBS (to achieve a final concentration of 1.2 mg iodine per gram of body weight) effectively delineated the primary tumor while minimizing hyperdense artifacts that can compromise both primary tumor and paraspinal muscle volumetric measurements on CT imaging^18^.

Our study also reports the novel application of Vivovist for the detection and volumetric tracking of liver metastases in preclinical cancer models. Vivovist, a nanoparticle-based contrast agent composed of alkaline earth metal cores, is primarily described in the literature for intravenous use to enhance vascular imaging^19^. Here, we demonstrate that a single intraperitoneal dose of Vivovist (1.5 g/kg) enabled reliable distinction between liver parenchyma and metastatic lesions for the duration of our study–up to 72 days post-injection. In contrast, hepatocyte-specific contrast agents such as 1,3-bis[7-(3-amino-2,4,6-triiodophenyl)heptanoyl]-2-oleoyl-glycerol have been shown to clear rapidly via the biliary system, limiting their window for effective tumor detection to within 24 hours post-injection^20,21^. This dual-contrast protocol is broadly adaptable to other preclinical models involving abdominal tumors or liver metastasis– common in many solid tumors, such as colorectal, pancreatic, and breast cancers^22^.

Our dual-contrast microCT protocol was particularly advantageous during early tumor development and metastatic seeding–critical timepoints when conventional imaging modalities are lacking. Ultrasound (US), while widely used, has limited sensitivity in detecting small, irregularly shaped tumors or liver and lung metastases, due to rib interference. Bioluminescence imaging (BLI), though convenient for whole-body surveys, can produce ambiguous signals when multiple organs overlap or when tumors reside near high-background tissues like the liver or gut. Additionally, commonly used reporters such as luciferase and GFP have been shown to elicit immune responses in immunocompetent hosts^23^. In contrast, our dual-contrast microCT protocol provided high-resolution, volumetric delineation of both primary and metastatic lesions, even at early stages of disease. This capability is especially important in pancreatic cancer, which is characterized by aggressive, early-onset metastatic spread^24^–often before the primary tumor becomes symptomatic or radiographically apparent^25,26^. As a result, many patients present with advanced disease, limiting therapeutic options and contributing to poor survival outcomes. Preclinical models that fail to capture early metastatic events are unlikely to recapitulate the clinical trajectory of human PDAC, and may overestimate the efficacy of investigational therapies. By enabling early visualization of metastases and their progression *in vivo*, our microCT protocol enhances mechanistic studies of metastatic seeding and organotropism and strengthens the translational value of preclinical therapeutic testing, particularly for agents targeting early dissemination or pre-metastatic niches.

Moreover, the capacity to longitudinally measure skeletal muscle volume in our study provided new insights into the kinetics of cachexia onset—an aspect of tumor biology that is often overlooked in preclinical models and is a major contributor to PDAC mortality^27^. We found that skeletal muscle loss occurred in a tumor-line-dependent manner and could be detected earlier than would be appreciated by endpoint measurements alone. This finding highlights an underrecognized heterogeneity in cachexia progression and underscores the importance of dynamic, temporal assessments. Further our approach is analogous to opportunistic use of clinical diagnostic CT scans to quantify muscle loss^28^, providing translatability to our platform.

While frequent imaging improved resolution and statistical power by reducing the number of animals required, we recognize that cumulative radiation exposure and imaging demands are non-trivial. However, control experiments support that radiation effects did not impact our study endpoints, such as primary tumor growth, metastasis burden, or cachexia. Future applications of this protocol may benefit from optimizing scan frequency based on experimental phase–e.g., increasing frequency during early metastatic seeding or cachexia onset, while decreasing frequency during stable tumor phases–to balance information yield with animal welfare and resource constraints.

Together, our findings establish dual-contrast microCT as a robust and versatile platform for preclinical cancer research. By enabling integrated assessment of primary tumor growth, metastatic dissemination to multiple organs, and systemic wasting over time, this imaging protocol expands the toolkit for studying complex cancer phenotypes and evaluating therapeutic interventions in preclinical models.

## Acknowledgements

This work was supported by grants from the Department of Veterans Affairs I01CX002046 (TAZ), the National Institutes of Health: P30 CA069533 to the OHSU Knight Cancer Institute, R01 CA257452 (TAZ), P01 CA236778 (TAZ), R01 CA287672 (JRB), R01 CA212600 (JRB), U01 CA224012 (RCS & JRB), R21 CA263996 (RCS & JRB), U01 CA294548 (RCS & JRB), U01 CA278923 (RCS), R01 CA186241 (RCS), and T32 GM141938 (KRP), and the Department of Defense: PA210068RCS (RCS), 0011775448-0001 (PJW), and PA210068-P1 (PJW). PJW received support from NCI 3P30CA069533-24W2 Early-stage Surgeon Scientist Program. Additional support was provided by the Brenden Colson Center for Pancreatic Care, and the Krista L. Lake Endowed Chair (RCS). We also thank Dr. Alexander Guimaraes for insightful discussions on radiological aspects of this study, members of the Zimmers lab for their valuable input, and Xiaoyan Wang for technical support.

## Materials and Methods

### Animals

Male C57BL6/J mice (JAX, catalog #000664), aged 11 weeks at surgery (PDAC models) and female CD2F1 mice (C26 model) were housed in pathogen-free conditions at 22.5°C and 12 h light/12 h dark cycles. Animals were provided ad libitum access to water and food (Rodent Diet 5001; Purina Mills). Animals were scanned at the designated days post-tumor injection from 0800–1100 h. Mice were monitored daily for health status and humane endpoints were defined by tumor weight ≥ 10% mouse body weight or a Hickman score ≤ 3, based on body condition score, behavior, and appearance^10^. All studies were conducted in accordance with the National Institutes of Health Guide for the Care and Use of Laboratory animals and approved by the Institutional Animal Care and Use Committee of Oregon Health and Science University.

### Tumor Models

All lines were tested for adventitious viruses (Charles Rivers Laboratory). Six murine PDAC cell lines were used: KPC 32908, KPC 7107, KPC 8060, KMC Z682, KMC Z693, and KMC Z696 (KPC– LSL-*Kras^G12D/+^;*LSL-*Tp53^R172H/+^;Pdx1-Cre*; KMC– LSL-*Kras^G12D/+^*;*Rosa26*- LSL*^Myc/Myc^*;*Ptf1a^CreERTM^*, updated from ^12^). The KPC 7107 and KPC 8060 lines were kindly provided by Dr. Tony Hollingsworth (University of Nebraska Medical Center) and the KPC 32908 line was kindly provided by Dr. David Tuveson (University of Pennsylvania; derived from ^11^). The KMC Z693 and Z696 lines were used in the study comparing primary tumor volume measurements derived from 2D ultrasound versus microCT (Fig. 5). All lines were cultured in DMEM supplemented with 10% fetal bovine serum and 1% penicillin-streptomycin. <u>Orthotopic</u> <u>PDAC model:</u> Immediately prior to surgery, mice were given a flank subcutaneous (SQ) injection of Meloxicam (5mg/kg). Under isoflurane anesthesia, a small surgical incision was made in the upper left quadrant of the abdomen, retracting the skin layers, fascia, and muscle wall to expose the pancreas. The tail of the pancreas was injected with 5,000 cancer cells suspended in 20 μl of 1:1 Matrigel and phosphate-buffered saline (PBS) or an equal volume of cell-free Matrigel and PBS. Post-wound closure, mice received a flank SQ injection of 400μl warm sterile PBS and the wound area was covered with 2.5% bupivacaine hydrochloride. Mice received a second flank SQ injection of Meloxicam (5mg/kg) 24 hours post-surgery. <u>Subcutaneous colon cancer</u> <u>model:</u> One million C26 tumor cells (National Cancer Institute) were injected subcutaneously in the right flank under isoflurane anesthesia.

### Radiation versus Ultrasound Study

To assess the effects of radiation on primary tumor growth, metastasis, and cachexia, 5,000 KMC Z682 PDAC cells were orthotopically injected into 20 male C57BL6/J mice (JAX, catalog #000664). Mice were assigned to either a non-radiation (N=10, ultrasound imaging) or radiation (N=10, microCT imaging) group. Mice were imaged starting 15 days post-injection, with scans performed weekly until day 29, then every four days until study endpoint (day 43). Time under anesthesia (isoflurane) and contrast administration (iohexol and Vivovist, following the below protocol) was matched between groups. At endpoint, all mice underwent microCT imaging for volumetric metastatic burden assessment immediately before euthanasia and tissue harvesting.

### MicroCT Contrast Administration

To visualize the primary tumor, iohexol (GE Healthcare, Omnipaque 300 mg I/mL) was diluted 1:3 in PBS and administered intraperitoneally (IP) at a final concentration of 1.2 mg iodine per gram of body weight immediately prior to microCT imaging. To enhance contrast between liver tissue and metastases, a single IP injection of 1.5 g/kg Vivovist (Nanoprobes #1301) was administered 24 hours before the first microCT scan.

### MicroCT Imaging and Reconstruction

Mice were imaged using a high-resolution SkyScan 1278 microCT scanner (Bruker) starting 7 days post-tumor cell injection and subsequently scanned every 3–4 days or as indicated. The scanner is equipped with a fully integrated physiological monitor, which tracks temperature, breathing, and ECG during the scan, alongside cumulative radiation dose. Prior to each day of scanning, the microCT machine was aged by turning on the radiation for approximately four minutes and then the flat-field correction was updated to ensure background is represented in a consistent grey level, controlling for interpixel intensity variations that would otherwise result in ring artifacts. Mice were scanned in the prone position to optimize paraspinal muscle measurements, as this orientation enhances muscle delineation from adjacent organs such as the kidneys. Scans were performed at a 50 μm pixel size, 0.5 mm Al filter, 40 ms exposure time, rotation step of 0.250°, and at 180° with no frame averaging. We gated our scans using the physiological monitoring window with thoracic movement as our trigger option. Image reconstruction was done with a 20% beam hardening correction and ring artefact reduction of 4 using CTrecon software (Bruker).

### Imaging Data Analysis

MicroCT reconstructions were analyzed using CTan and CTvol software (Bruker) to quantify primary tumor volume, metastatic burden, and paraspinal muscle area, as well as to generate 3D renderings. Primary tumors were segmented, and their volumes were extracted using CTan. For liver and lung metastases, each organ was manually scanned to identify individual lesions, and total metastatic burden was calculated by summing the volumes of all metastases. Given the predominantly spherical nature of these lesions, metastasis volume was approximated using the diameter of the largest cross-sectional area:

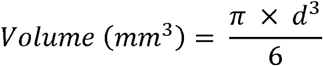

Paraspinal muscle area was measured at the most rostral cross-section of the L5 vertebra and normalized to baseline values. Endpoint tumor and muscle weights were recorded post-mortem to validate imaging-based measurements.

For ultrasound imaging, primary tumor volumes were estimated using the largest axial and sagittal cross-sectional areas, applying the following equations:

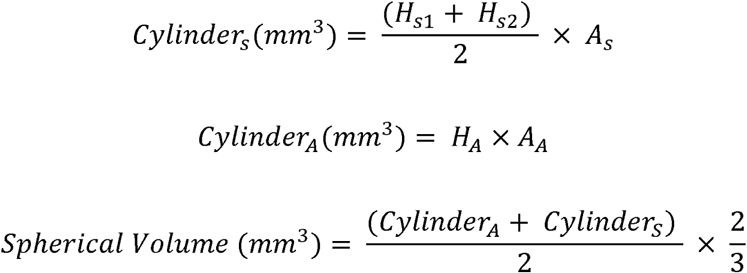

*H = height*

*A = area*

*X_A_ = X_axial_*

*^X^s ^= X^sagittal*

### Histological Analysis

Tissues were harvested at endpoint, fixed in 10% formalin for 24 hours at 4 °C, and embedded in paraffin for histology. Hematoxylin and eosin (H&E) staining was used for tumor and metastasis characterization. Immunohistochemistry was performed using HuR antibody (Proteintech, cat. # 11910-1-AP, rabbit, 1:400) and either DAB-conjugated (Cell Signaling Technology, cat. #8114, goat anti-rabbit) or alkaline-phosphatase-conjugated (Vector Laboratories, cat. #MP-5401, horse anti-rabbit) secondary antibodies.

### Statistical Analysis and Reproducibility

Statistical analyses were performed using GraphPad Prism 9. Tissue weights between lines and longitudinal paraspinal muscle % areas were analyzed using one-way ANOVA with Dunnett’s multiple comparisons test. Kaplan-Meier analysis with log-rank tests was used to assess time-to-metastasis and survival endpoints. Correlations between imaging-derived tumor volumes and endpoint tumor weights were assessed using Pearson’s correlation. Significance was defined as p < 0.05. In all studies, data represent biological replicates (n) and are presented as the meanLJ±LJstandard deviation.

---

**Supplementary Table 1.**
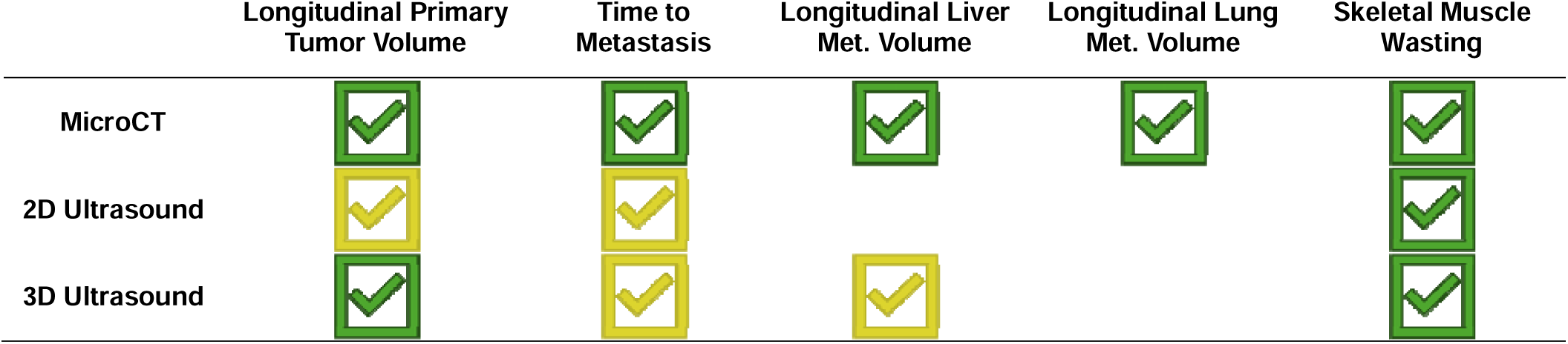
Imaging modality recommendations based upon primary endpoint. Green check means imaging modality is appropriate for endpoint. Yellow check means imaging modality is possible but not ideal for endpoint. Met = metastasis

**Supplementary Table 2.**
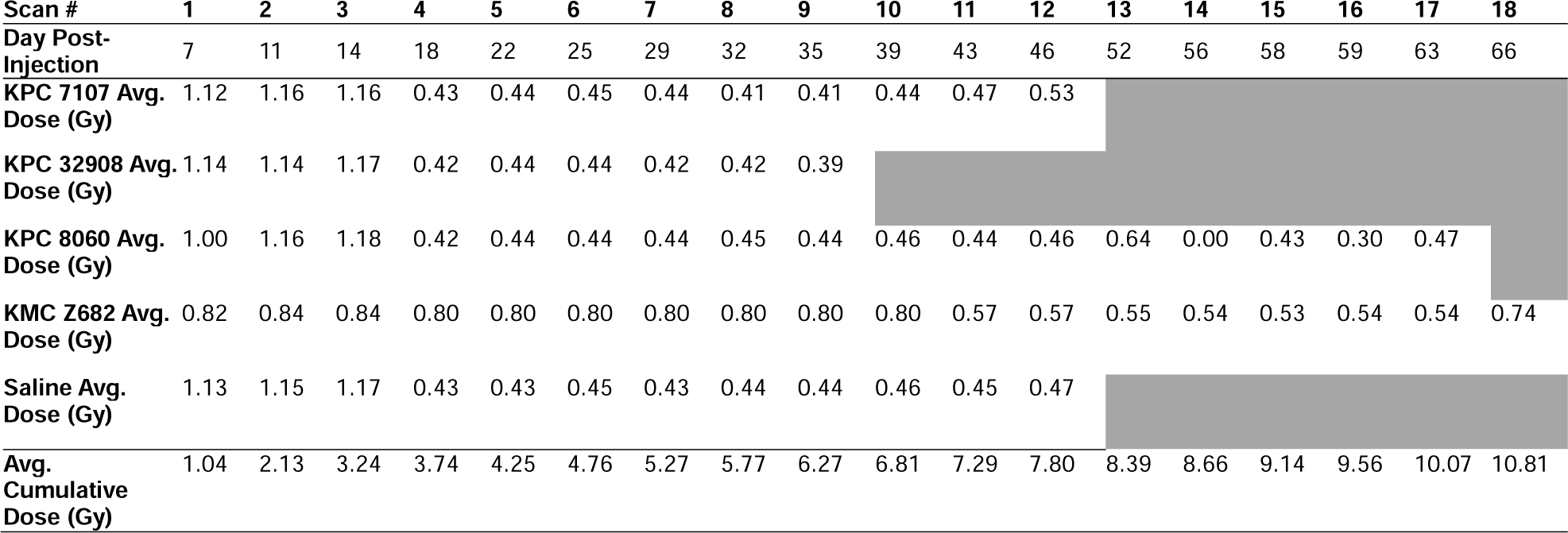
Cumulative radiation dose in reported study. . Discrepancies in average scan radiation dose are due to optimizing settings and gating throughout the study. KMC Z682 group was scanned at a separate time, which is why average radiation dose varies compared to the other four groups.

**Supplementary Fig. 1.**
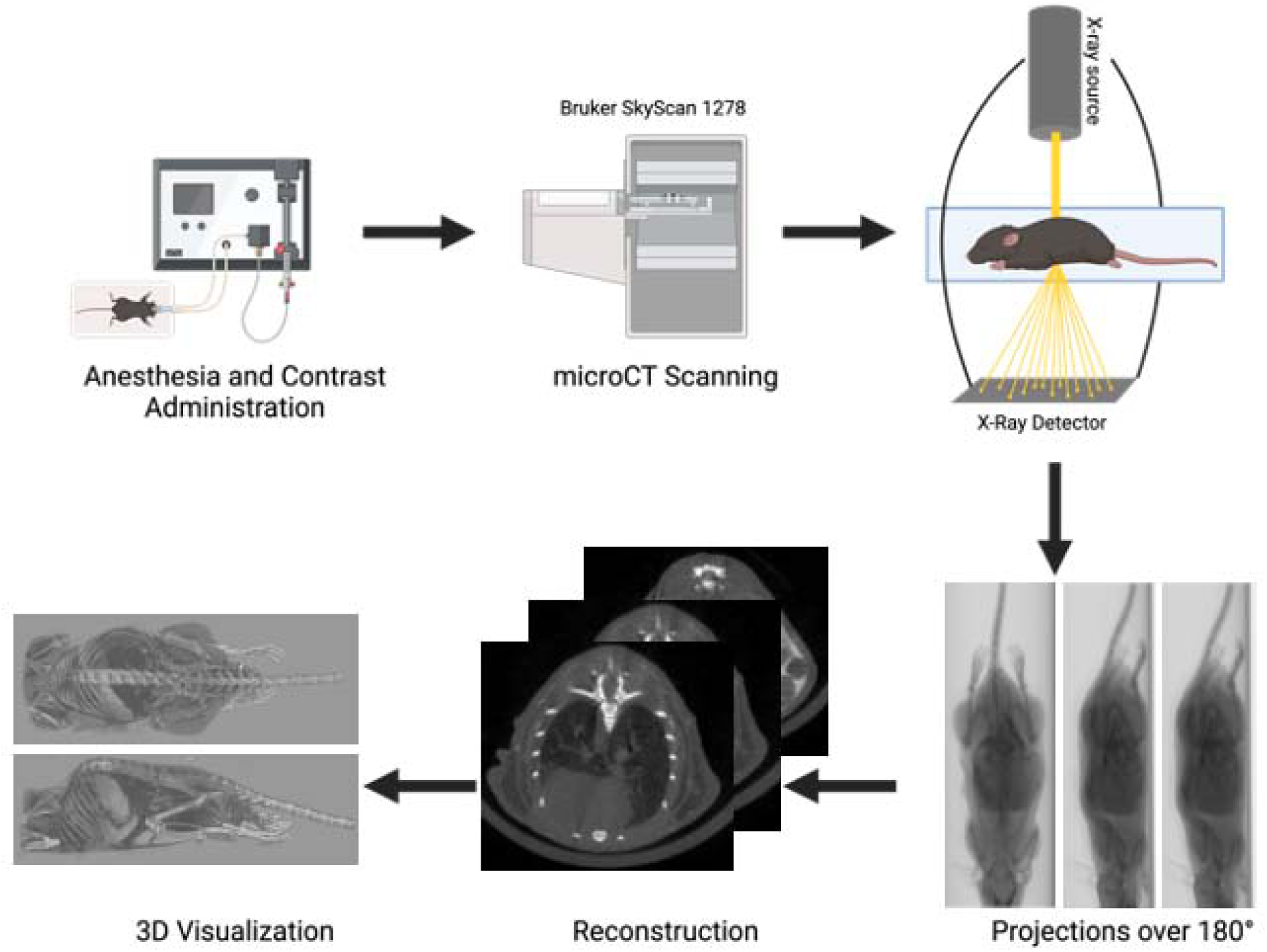
MicroCT workflow. MicroCT generates high-resolution, three-dimensional (3D) tomographic images by capturing hundreds of two-dimensional (2D) cone-beam projections around the mouse. The raw projection data are pre-processed using dark current and flat-field corrections before being reconstructed with the Feldkamp algorithm modified for cone beam using a Hamming window filter, incorporating geometric parameters to minimize misalignment artifacts. The resulting tomographic images consist of isotropic voxels, where voxel intensity reflects the mean linear attenuation coefficient, enabling visualization in multiple orientations as 2D slices or a rendered 3D volume^9^.

**Supplementary Fig. 2.**
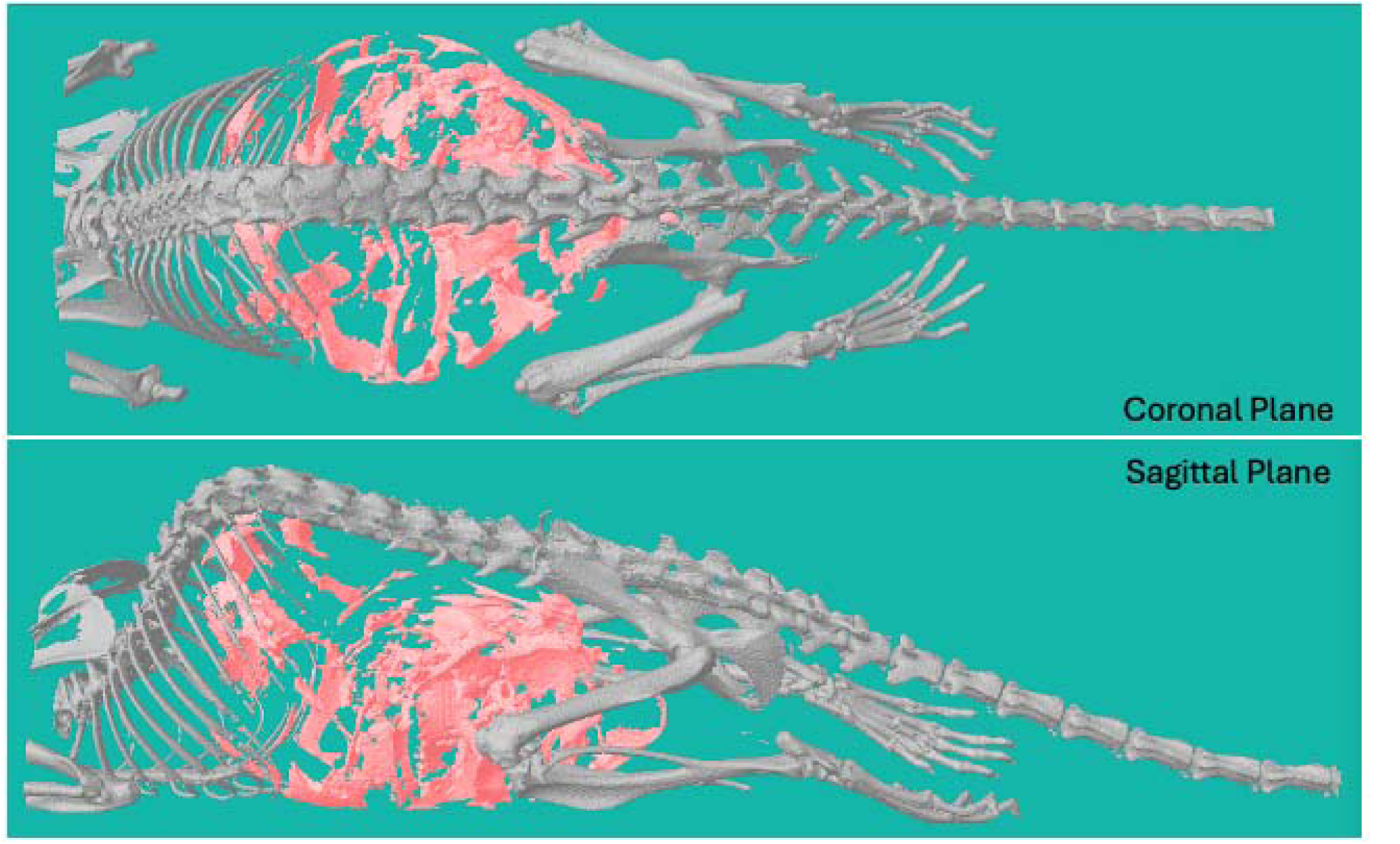
Biodistribution of iohexol contrast injected intraperitoneally. Iohexol (Omnipaque, 300 mg I/mL) was diluted 1:3 in PBS (final concentration of 1.2 mg iodine per gram of body weight) and administered intraperitoneally immediately prior to microCT scanning. The figure shows a representative 3D reconstruction in coronal (top) and sagittal (bottom) planes, with iohexol contrast highlighted in red. The skeleton of the mouse is rendered in grayscale, illustrating the distribution of the contrast medium surrounding organs within the open space of the peritoneal cavity.

**Supplementary Fig. 3.**
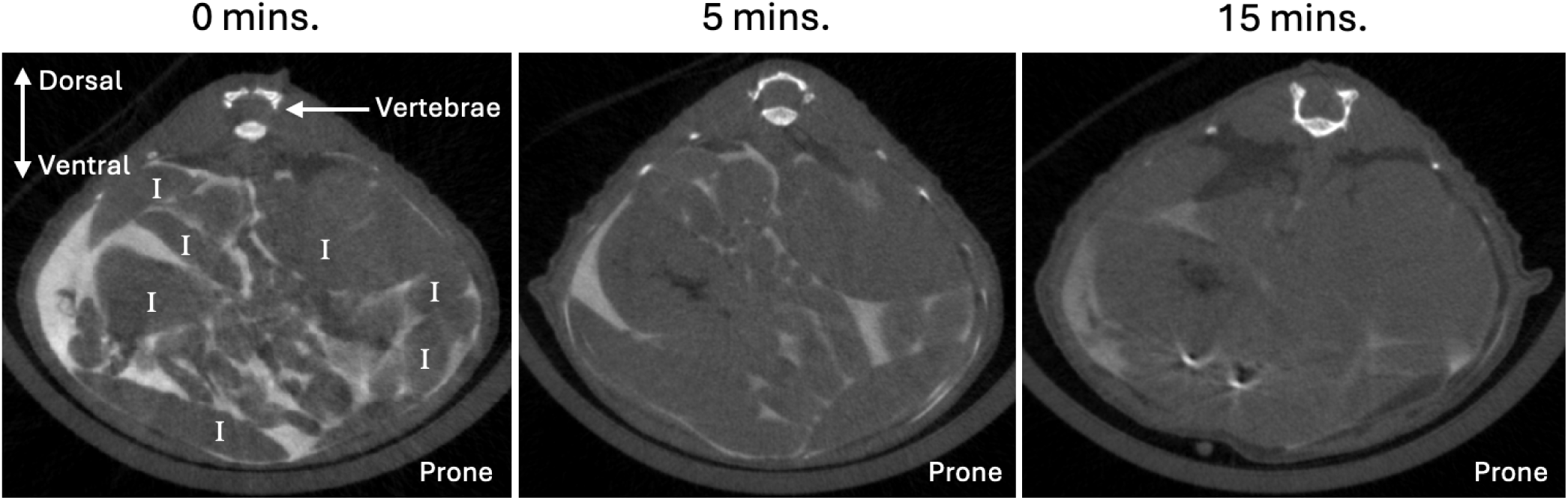
Time course of iohexol contrast distribution following intraperitoneal injection visualized by microCT imaging. Mice were injected with iohexol contrast (Omnipaque, 300 mg I/mL, diluted 1:3 in PBS, final concentration 1.2 mg I/g) intraperitoneally and scanned at 0-, 5-, and 15-minutes post-injection. Representative axial microCT images show contrast initially distributed throughout the peritoneal cavity (0 min), progressively diffusing by 5 minutes, and largely clearing from the peritoneal space by 15 minutes. By 20 minutes post-injection, contrast was predominantly localized in the bladder (data not shown). I = intestine.

**Supplementary Fig. 4.**
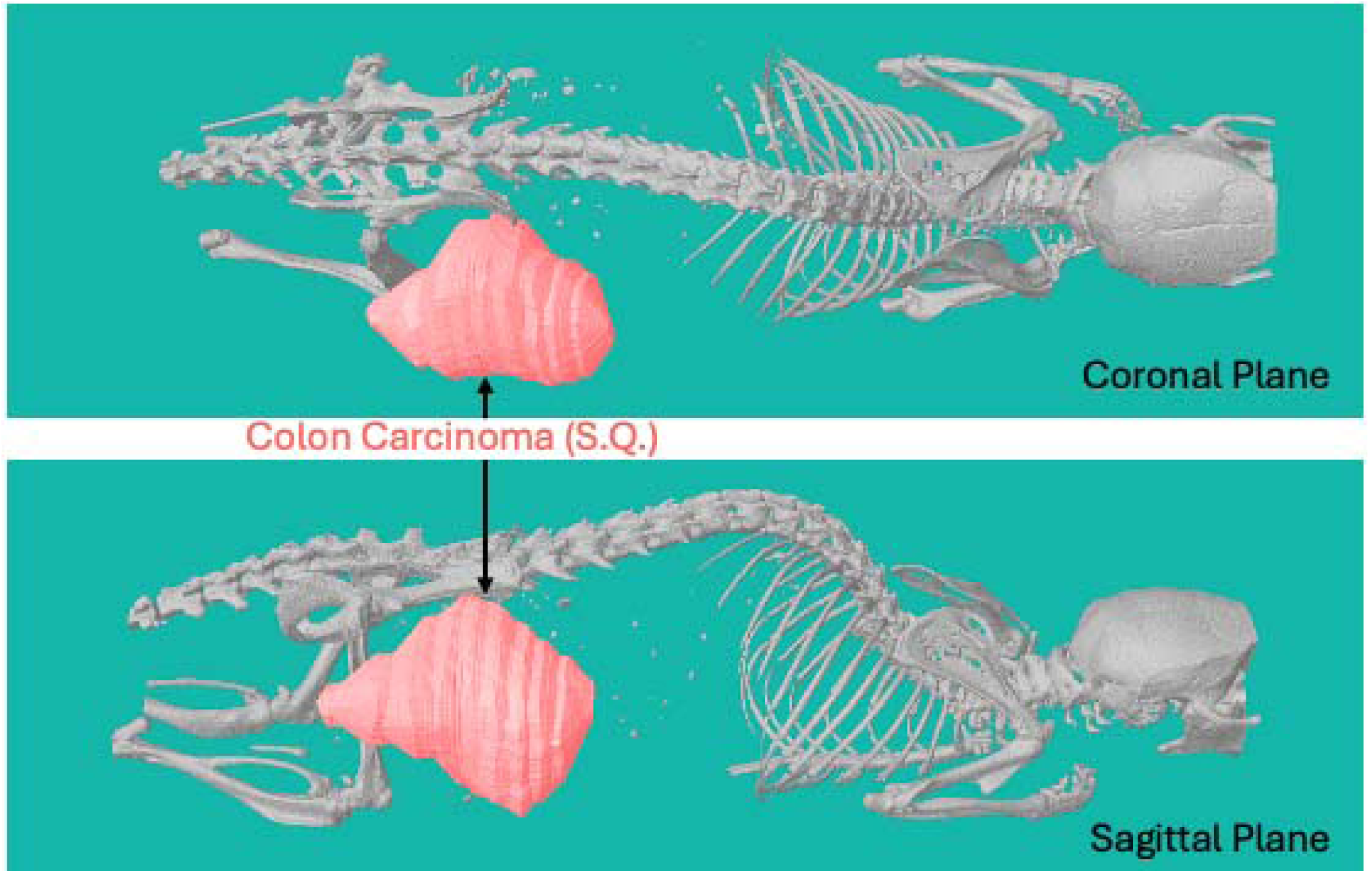
MicroCT 3D reconstruction of subcutaneous tumor from C26 colon cancer model. Representative 3D reconstruction in coronal (top) and sagittal (bottom) planes, showing a subcutaneously injected C26 colon carcinoma tumor (highlighted in red). The skeleton of the mouse is rendered in grayscale. S.Q. = subcutaneous.

**Supplementary Fig. 5.**
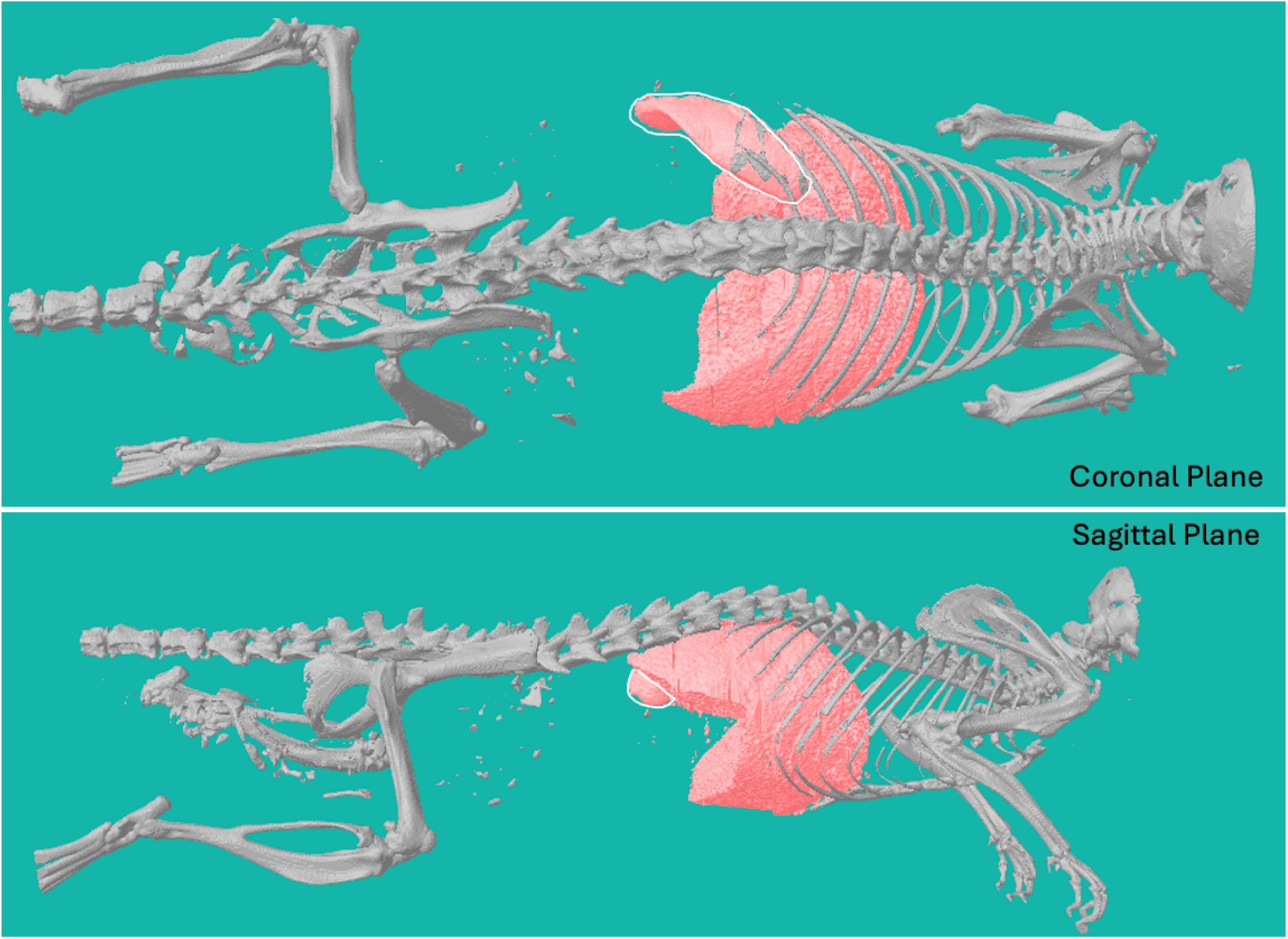
Biodistribution of Vivovist contrast. A single intraperitoneal injection of 1.5 g/kg Vivovist (Nanoprobes #1301) was administered 24 hours before the first scan. Vivovist, composed of alkaline earth metal nanoparticles, is rapidly taken up by the liver and spleen, maintaining consistent contrast intensity for the study duration (up to 72 days post-injection). Representative 3D reconstruction in coronal (top) and sagittal (bottom) planes from a scan on day 11 post-contrast injection, with Vivovist contrast highlighted in red. The skeleton of the mouse is rendered in grayscale, illustrating the distribution of the contrast medium within the spleen and liver. Spleen is outlined in white.

**Supplementary Fig. 6.**
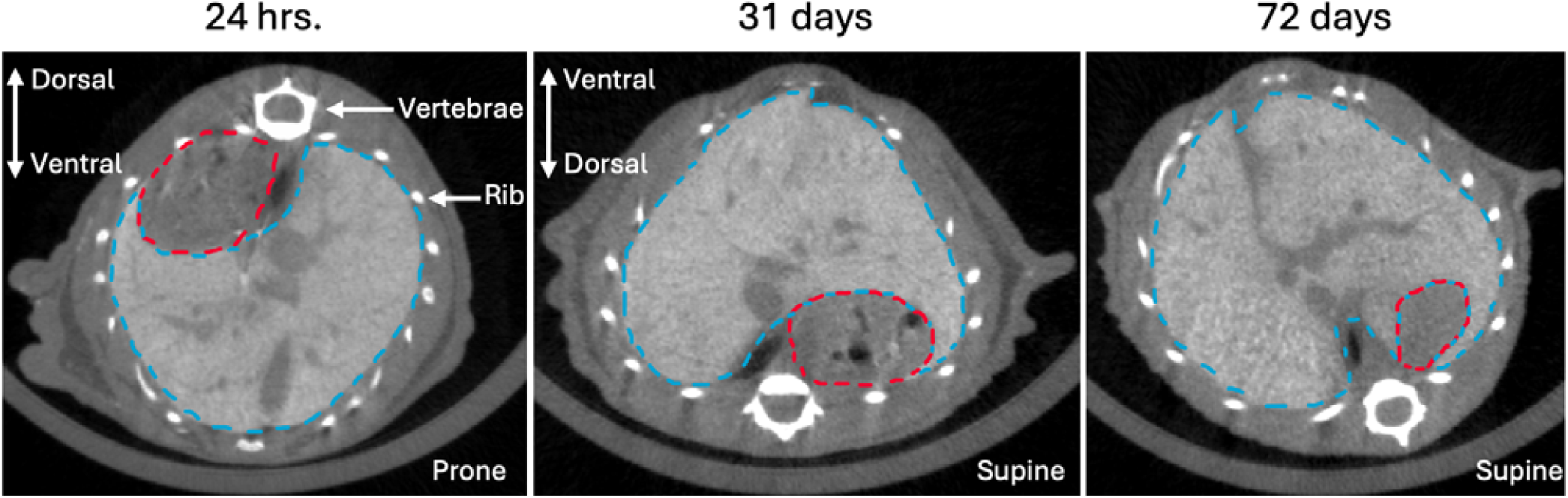
Longitudinal retention of Vivovist contrast visualized by microCT imaging. Representative axial microCT images of a mouse 24 hours, 31 days, and 72 days after intraperitoneal injection of Vivovist (1.5 g/kg, Nanoprobes #1301) contrast agent. Contrast retention in the liver is evident, with persistent signal intensity even at 72 days post-injection. Mice were scanned in the prone position at 24 hours and in the supine position for subsequent time points. Stomach (outlined in red); Liver (outlined in blue).

**Supplementary Fig. 7.**
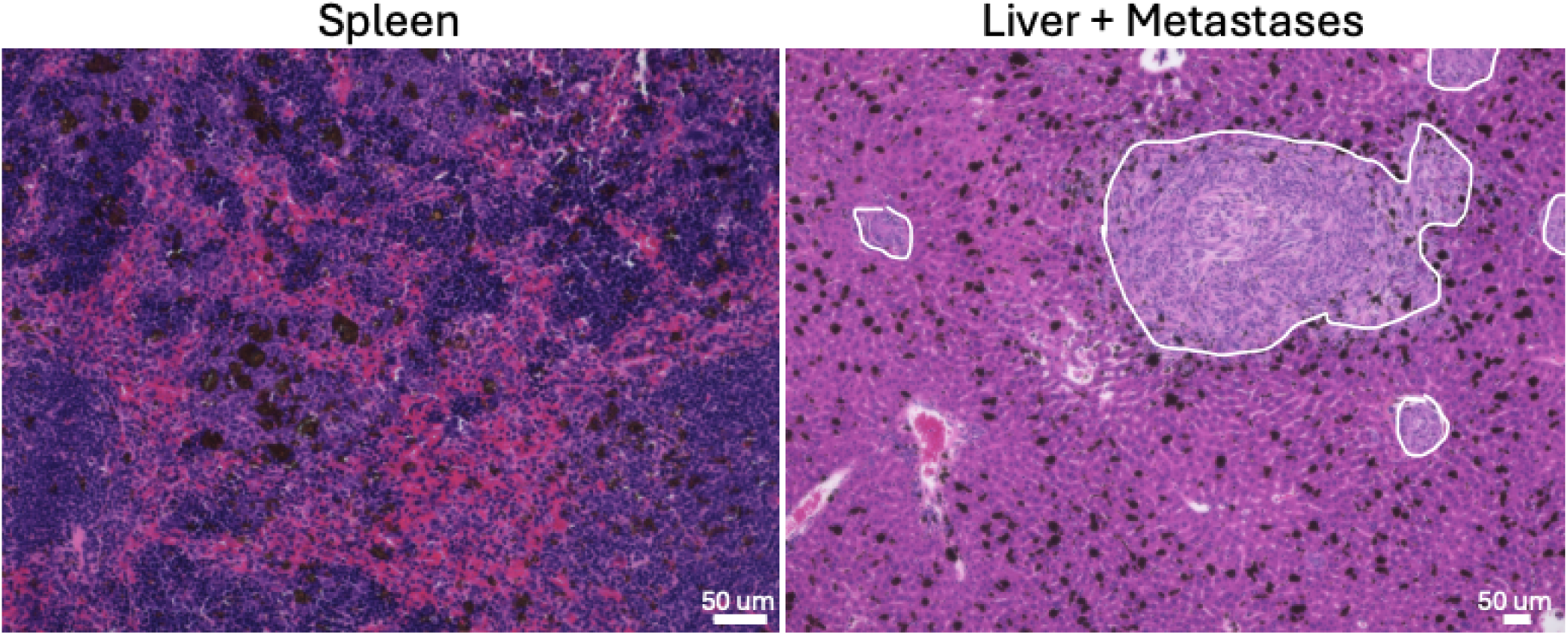
Vivovist biodistribution visualized by hematoxylin and eosin (H&E) staining. Representative H&E-stained sections showing Vivovist uptake in the spleen (left) and liver (right). Vivovist does not get taken up in the liver metastases (metastatic lesions in the liver are outlined in white), effectively distinguishing liver metastases from normal tissue. Vivovist particles appear as dark brown deposits. Scale bars: 50 µm. Spleen (left) and liver (right) images taken at 20X and 10X, respectively.

**Supplementary Fig. 8.**
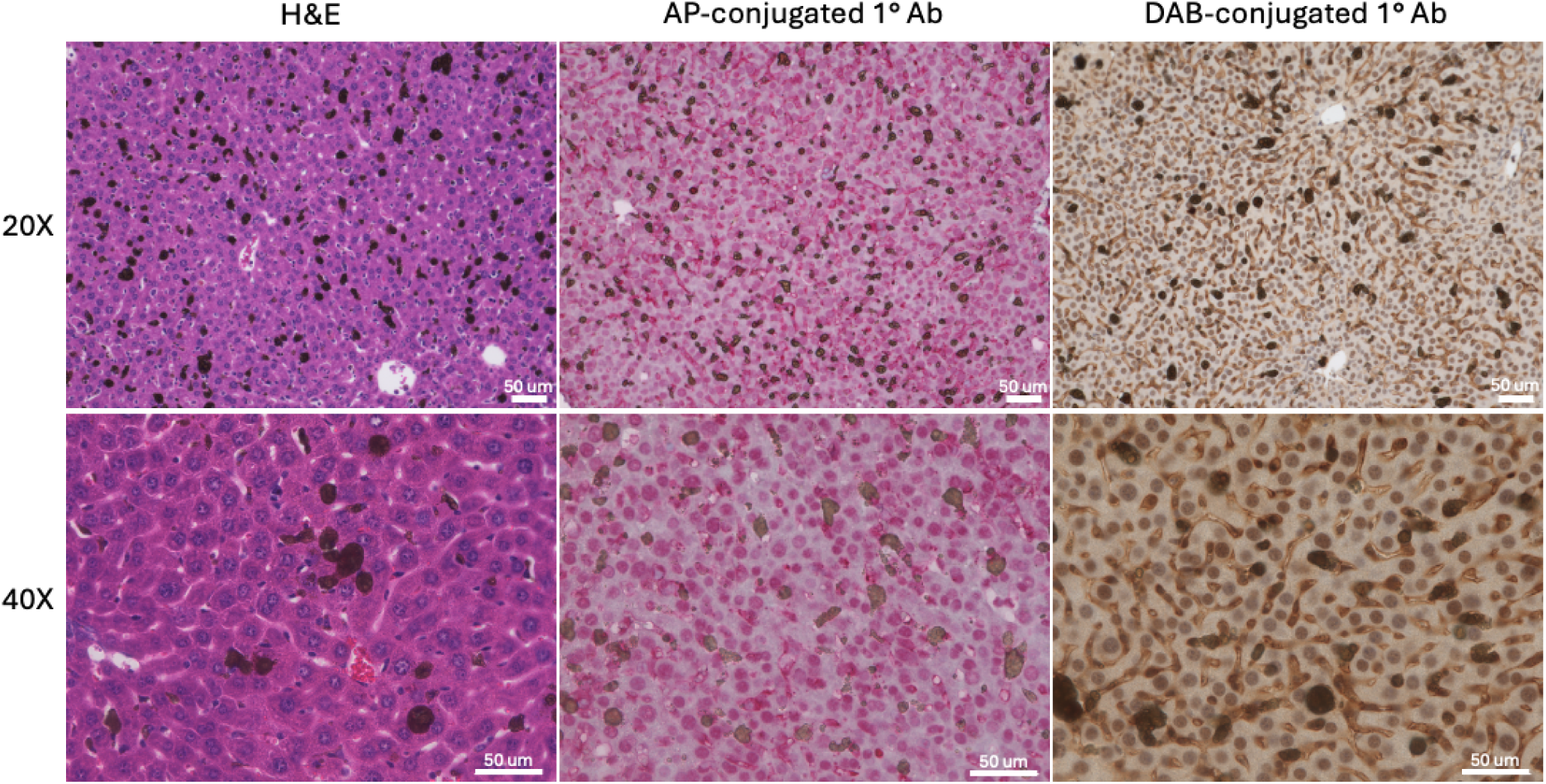
Vivovist contrast artifacts disrupt immunohistochemistry visualization and strategies to mitigate interference. Hematoxylin and eosin (H&E) (left) and immunohistochemistry (IHC) using antibodies against HuR, which localizes to both the nucleus and cytoplasm, conjugated to alkaline phosphatase (AP, middle) or 3,3′-diaminobenzidine (DAB, right) demonstrate a persistent dark brown hue in liver tissue due to retained Vivovist contrast agent. This artifact can obscure brown DAB IHC signal, complicating protein quantification. Use of AP-conjugated antibodies, which produce red staining, or switching to immunofluorescence is suggested to mitigate contrast artifact signal interference. Scale bars: 50 µm. Images taken at 20X (top row) and 40X (bottom row).

**Supplementary Fig. 9.**
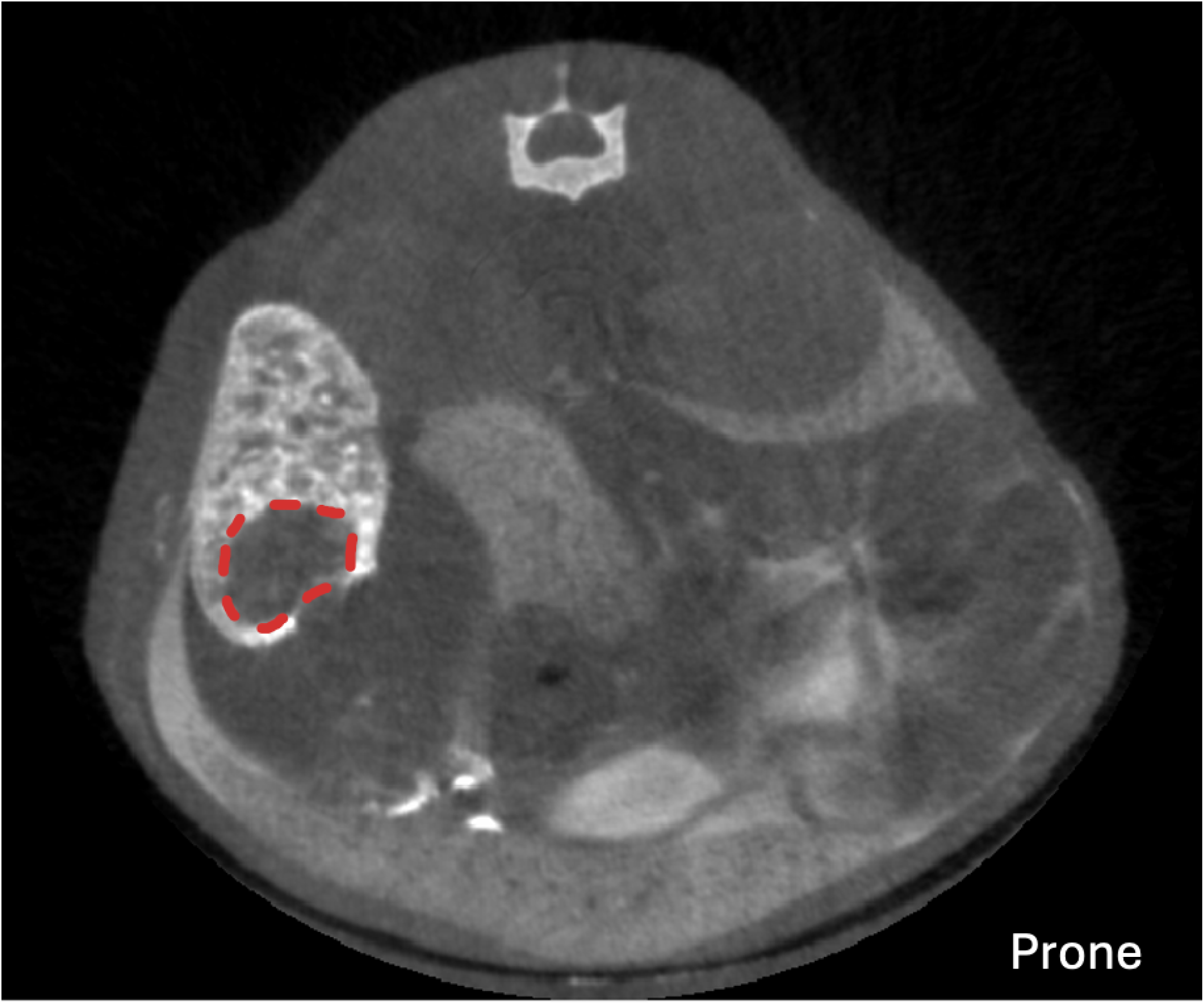
MicroCT slice demonstrating a metastasis in the spleen. Representative axial microCT image showing metastasis (outline with dashed red line) in spleen. Spleen is bright due to uptake of Vivovist contrast.

**Supplementary Fig. 10.**
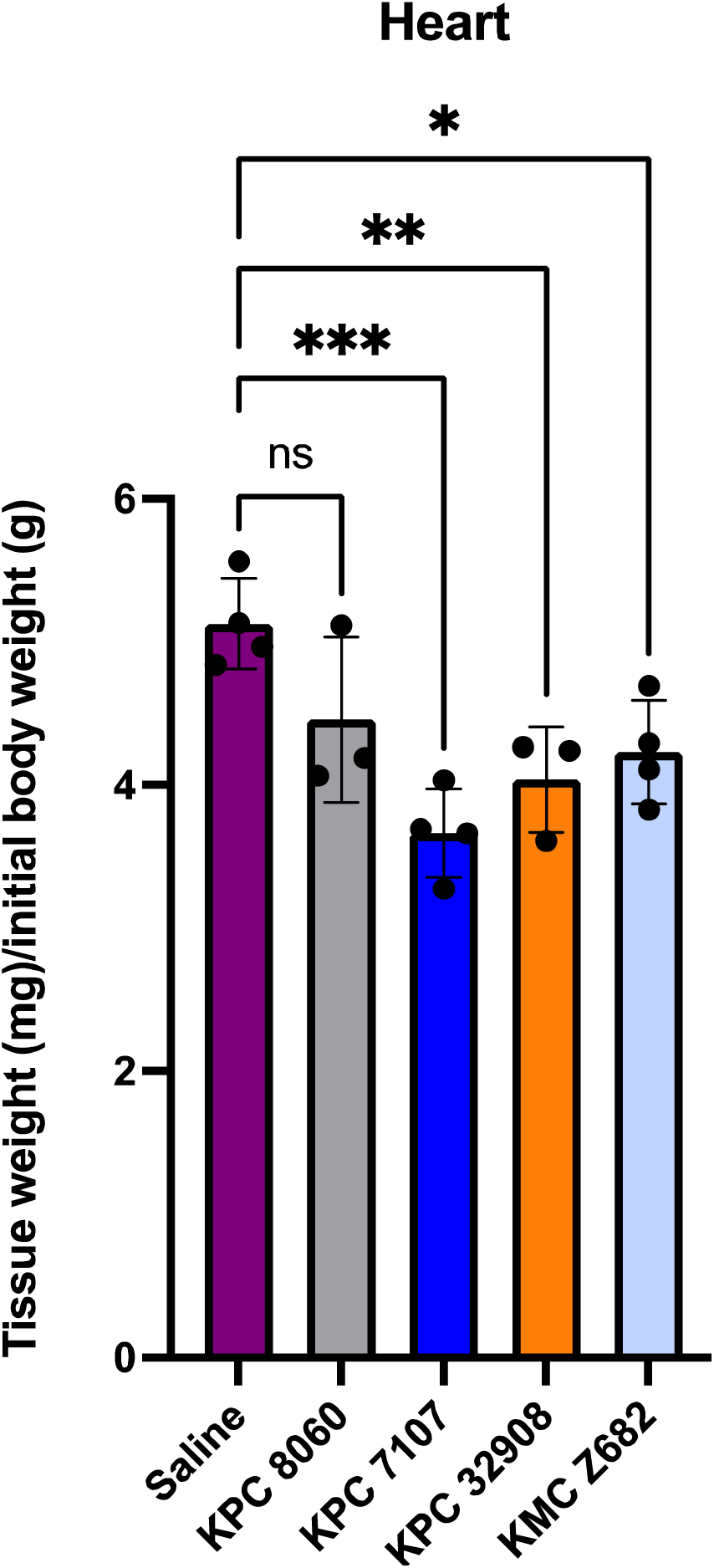
Tumor-induced wasting of heart tissue. Post-mortem analysis of heart weight normalized to initial body weight across experimental groups (N = 4 per group). Statistical analysis used one-way ANOVA with Dunnett’s multiple comparisons test. ns = not significant, * p < 0.05, ** p < 0.01, *** p < 0.001. Error bars represent mean ± standard deviation.

**Supplementary Fig. 11.**
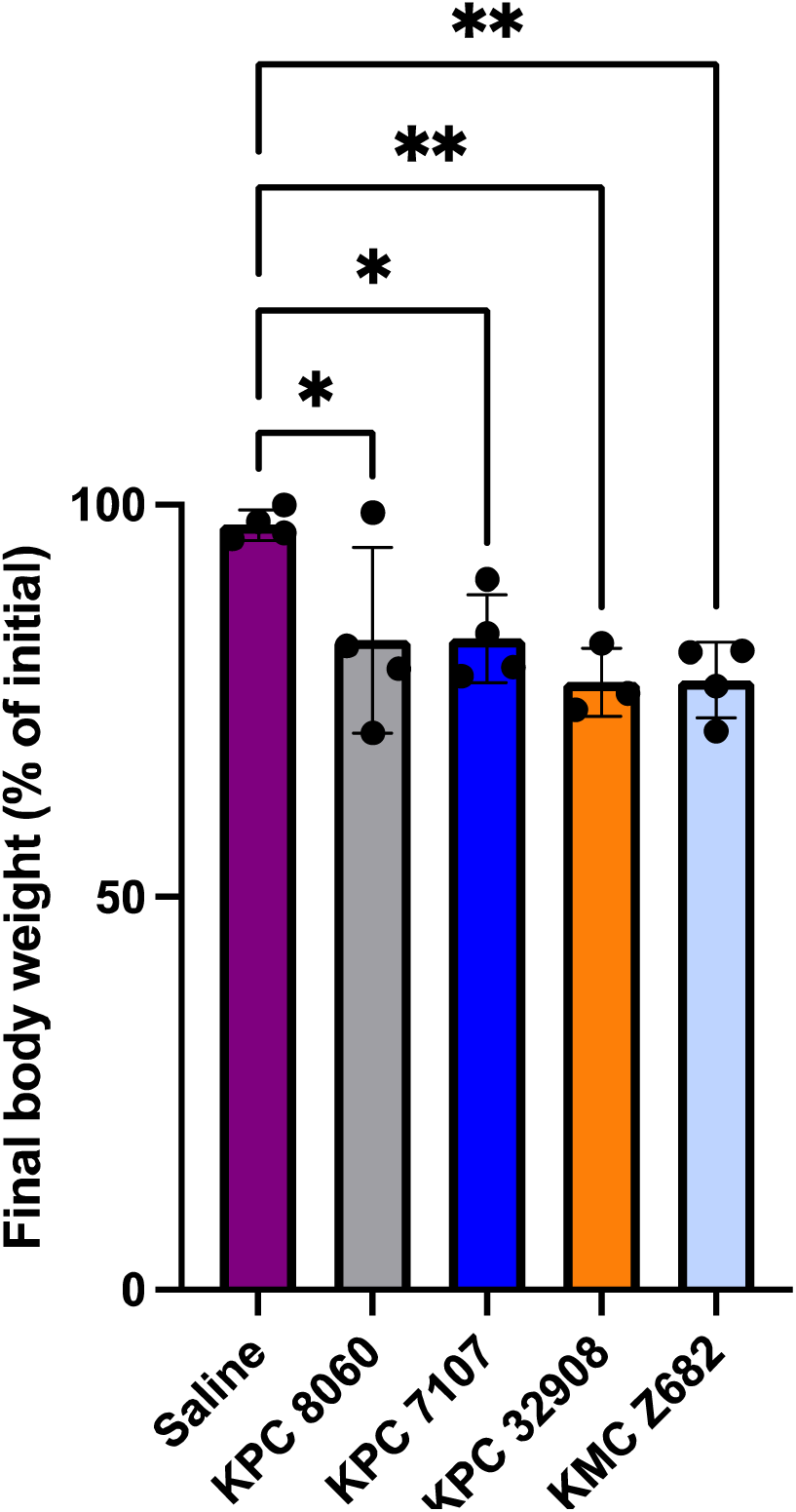
Tumor-induced weight loss is independent of radiation dose. Final body weight excluding tumor weight as a percentage of initial body weight for mice injected with different PDAC lines (KPC 7107, KPC 32908, KPC 8060, and KMC Z682) or PBS control (N = 4 per group). While all groups received comparable doses of radiation, only the PDAC tumor-bearing groups exhibited significant weight loss, suggesting that weight loss is driven by tumor burden rather than radiation exposure. Statistical analysis was performed using one-way ANOVA with Dunnett’s multiple comparisons test. * p < 0.05, ** p < 0.01. Error bars represent mean ± standard deviation.

